# The transmission of African Animal Trypanosomiasis (Nagana) in two districts in the forest zone of Ghana

**DOI:** 10.1101/2021.08.11.455898

**Authors:** Austine Tweneboah, Jana Rosenau, Addo Kofi Agyapong, Thomas Kwame Addison, Mahamat Alhadj Moussa Ibrahim, Judith Sophie Weber, Soerge Kelm, Kingsley Badu

## Abstract

The African Animal Trypanosomiasis (nagana) is caused by several species of trypanosome species which results in significant clinical diseases. This consequently leads to huge economic losses. We carried out a cross-sectional survey to investigate the composition of vectors, parasite diversity and intensities in two districts in the Eastern region of Ghana. Cytochrome c oxidase subunit 1 (COI) and internal transcribed spacer (ITS1) dependent PCR assays identified tsetse fly species and trypanosome parasites circulating in the area. A total of 229 tsetse flies, 65 pigs and 20 were sampled. Female and male flies 155 (51.8%), 74 (32.3%) respectively were collected with biconical traps. The overall vector density of 4.3 flies/trap/day was observed. Typanosome prevalence of 58.9% (95% CI: 52.5–65.1), 46.2% (95% CI: 34.6–58.1) and 0.0% (95% CI: 0.0–16.1) in tsetse flies, pigs and cattle respectively were detected. *Trypanosoma congolense* was the predominant species with a prevalence of 80.7% (95% CI: 73.3–86.5) in flies and 60.0% (95% CI: 42.3–75.4). There was evidence of multiple trypanosome infection with *T. congolense*/*T. simiae* occurring highest with prevalence of 38.0% (95% CI: 30.7–46.9). The parasite prevalence in pigs across the communities was high with significant differences associated between locations (Chi^2^ = 28.06, 95% CI: 0.05–0.81, P=0.0009). Blood meal with flies revealed feeding on both domestic *Sus scrofa domesticus* (pig) and *Phacochoerus africanus* (warthog). Tsetse flies in this area remain highly infected with trypanosomes and continue to transmit the parasites to livestock and human populations in the communities.

## Introduction

Trypanosomiasis is a vector-borne infectious disease of huge public health and economic importance in sub-Saharan Africa [1–3]. It is a vector-borne parasitic disease caused by a flagellate protozoan belonging to the genus *Trypanosoma* (T). These parasites circulate in the blood and central nervous system of their host (humans and livestock) and are transmitted by bloodsucking dipterans of the genus *Glossina* [4]. Several parasite species cause infection in humans and animals. *Trypanosoma brucei gambiense* [4] determines the human’s chronic disease form in West and Central Africa while *Trypanosoma brucei rhodesiense* [5] circulating in East and Southern Africa determines the acute form of the disease. The World Health Organization (WHO) estimates suggest 65 million people being at risk of Trypanosomiasis with a high proportion going without any medical monitoring. The disease in animals though is known as the African Animal Trypanosomiasis, otherwise known as ‘nagana’, a Zulu-originated word which means ‘powerless’ or ‘useless’ [6]. The parasite species infecting animals include *T. congolense, T. vivax, T. brucei. brucei, T. simiae* and several other subspecies. Unlike in domestic animals, where severe infection usually leads to death if untreated, the disease has relatively mild effects on wild animals [7–9]. At least 17,500 new cases are reported every year in meat production areas.

The disease is mostly restricted to rural forested communities that rely heavily on agriculture [5, 6]. As reported by WHO in 2010, the African continent is known to account for 90% of total number of reported cases for Human African Trypanosomiasis (HAT). In addition, nagana accounts for huge losses in animal production, generating loss of nearly $ 4.5 billion each year [10,5],13, 14]. As a result, Africa loses up to US$1.5 billion annually [15]. Information on the current situation of HAT in Ghana remains scanty as reported by WHO [16]; however, ‘nagana’ is mostly transmitted by *Glossina palpalis palpalis* which is the most widely distributed tsetse fly species in Africa, as such it remains highly endemic in Ghana[16–18].

The presence of tsetse flies is a threat to livestock production [20] in the Suhum municipality as many pigs have suffered death in the past (personal communication with Mr. Gbeve, Veterinary Officer). This study was therefore carried out to determine the prevalence of trypanosome parasites in the pigs and cattle blood as well as the invertebrate host (vector) using molecular techniques to identify disease risk areas in the Suhum municipality for future studies.

## Methodology

### Study area

This study was conducted in the selected communities in the Ayensuano and Kraboa districts located 6° 2′ 3.84″ N, 0° 27′ 8.64″ W in the Eastern region of Ghana. The region is positioned in the semi-deciduous forest zone with an annual rainfall of 1500 mm. Thus, most of the communities at the outskirts of the Suhum municipality have more than half of their population engaged in agriculture. The type of soil in the region is conducive for the cultivation of cash and food crops. As a result, crop farming and rearing of livestock are the main agricultural activities in the district. Farmers in remote areas have satellite homes with a small domestic farming as supporting income or the main economic livelihood for the family. Two different areas within the municipality were visited in February 2018 and February 2019. Previous studies indicated no variation in transmission across the climatic seasons [21]

### Study design

A cross-sectional community-based survey was employed where all pigs, cattle and human participants were sampled within the study period.

### Fly trapping and dissection

The flies were captured using un-baited biconical traps [8, 21–23] that were mounted close to the pigsties but under a shade to minimise fly mortality. The coordinates of each trap were recorded with a Global Positioning System. After fly collection, an index of apparent abundance (IAA) of tsetse was calculated as the number of flies found per trap per trapping day [25]. The flies were grouped into males and females based on the presence or absence of the hypopygium (a round structure on the last segment of the abdomen, figure 2) [26]. Each fly was immobilized at 4° C prior to dissection, and the wings, legs, proboscis, and midgut each placed into 2 ml micro centrifuge tubes for preservation. However, only the midgut was investigated for trypanosomes in this study.

**Figure 1.**
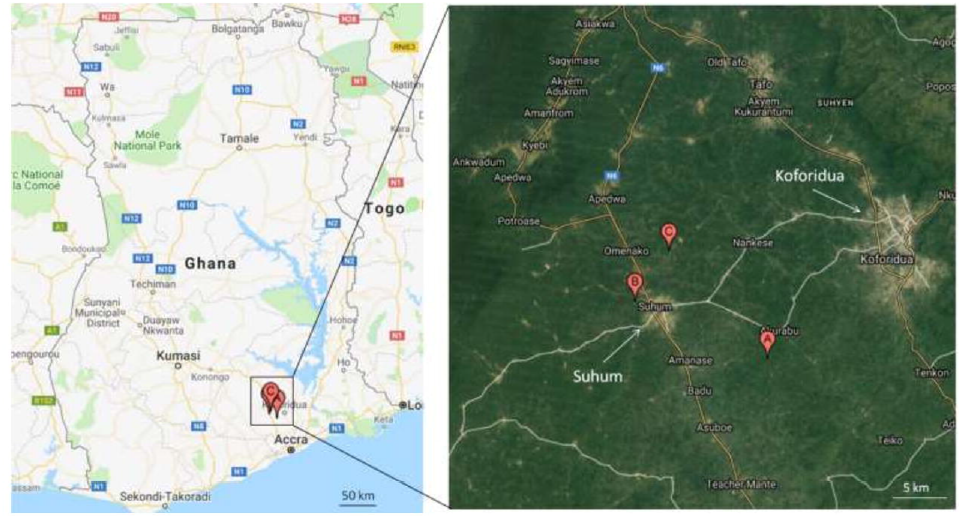
Tsetse flies were collected in some selected communities in the Suhum/Kraboa/ Coaltar district. Two traps were set up at Santramor (A), two at Pinto (B) and two at Adenta (C), one at Nkatekwan, and one at Ayekokooso.

**Figure 2:**
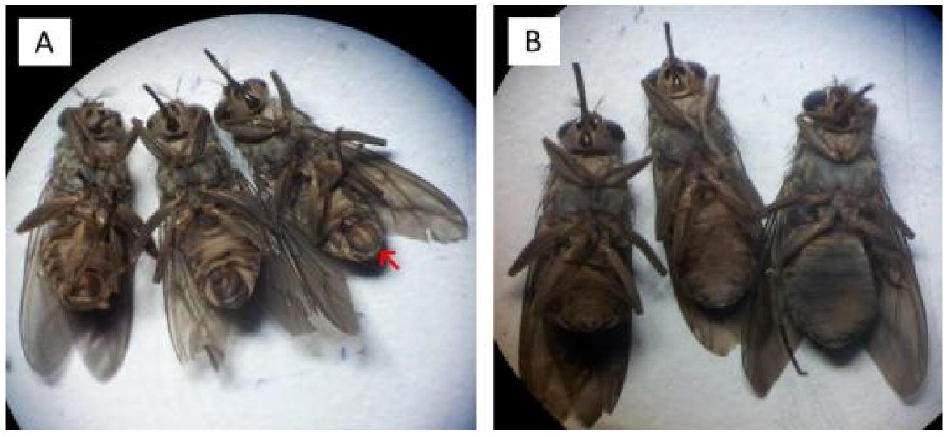
**A**: Male tsetse flies trapped in the field were identified by the presence of a hypopygium (at the external genitalia indicated by the red arrow); **B**. Female tsetse flies were identified by the absence of a hypopygium on the last segment of the abdomen [31]

### Ethical consideration

Ethical clearance was obtained from the Committee on Human Research, Publications and Ethics KNUST and the Komfo Anokye Teaching Hospital. Informed consent was obtained from the farmers prior to sampling.

### Blood sample collection

Blood samples were collected from 20 cattle and 65 pigs. Each cow was bled for 6 mL of blood from the jugular vein into a heparinized vacutainer and same volume of blood was drawn from the marginal ear vein of each pig into an EDTA tube. After sampling, the vacutainers were kept in a refrigerator until further processing.

### DNA Extraction

#### Tsetse fly

The tsetse fly midgut was homogenized in Tris-buffer (50 mM pH 9.0) using a motor-driven homogenizer and micro pestles (Kimble® Kontes disposable pellet pestles, Sigma-Aldrich), and 100µL of each homogenate was transferred into 1.5mL Eppendorf tube. DNA was extracted from the midgut with DNeasy Blood and Tissue Kit (Qiagen) according to the manufacturer’s instruction and 60 µL of each was eluted and stored at -20 °C.

#### Cattle and pig blood

DNA was extracted from, pig and cattle whole blood using DNeasy blood and tissue kit in accordance with the manufacturer’s protocol. The extracted DNA was quantified using Nanodrop ND-1000 spectrophotometer (Thermo Fisher Scientific) at 260 nm.

### PCR Amplification of the ITS1 gene for the Identification of Trypanosome sp

The ITS1 region was amplified by nested PCR to identify trypanosomes in tsetse midgut, cattle and pigs using the primers [26, 27] listed in Table 1. Each PCR reaction was carried out in a total volume of 25µL mixture each containing 2µM of each primer, 0.2mM of mix dNTPs, 1X of Buffer (Dream Taq), 0.1U of Taq polymerase and molecular-graded water (All reagents from Thermo Fisher Scientific). The cycling conditions were the same for both nested reactions with denaturation at 94 °C for 1 minute, annealing at 54 ^0^C for 30 seconds, extension at 72 °C for 30 seconds. The reaction mixture was cycled for 30 times in each PCR. The mixture was incubated at 95 °C for 3 minutes prior to the cycling and there was final extension at 72 °C for 5 minutes after the cycling. All PCR reactions were carried out in a Mastercycler Personal (Eppendorf). The resulting PCR products were electrophoresed on a 1.5% agarose gel run in Tris-borate-EDTA buffer at 100V for 1 hour.

### *T. congolense*-specific PCR

Samples which showed a band-size of approximately 700bp with generic primers were further run with *T. congolense* specific primers to confirm the presence of *T. congolense*. The species-specific primers (Table 1) used in a nested PCR also targeted the ITS region of the parasite’s DNA. Sample preparation and PCR reactions were identical to ITS1 nested PCR as already described in this paper.

### Amplification of the COI gene for the identification of tsetse fly species

To identify the tsetse fly species in the study area, a part of the COI gene [29] was amplified in a one-step PCR following the protocol as described by Ngomtcho and others [28].

### Blood meal Analysis

The extracted DNA from tsetse midgut were screened for the mitochondrial cytochrome b oxidase (Cyt b) gene for all vertebrates in a one-step conventional PCR as described by Muturi and others [30].

Table 1. Primers, annealing temperature and target as used in this study

### Sequencing of COI fragments

COI fragments were sequenced for the identification of tsetse fly species in the study area. The resulting gel product from *cox1* PCR were purified using GeneJET Gel Extraction Kit (Cat. no. K0701, Thermo Scientific) according to the manufacture’s protocol. The gel containing the DNA fragments were excised with a clean scalpel into pre-weighed 1.5ml Eppendorf tubes. The cut gels were purified according to the manufacturer’s protocol and 50µL of each sample was collected in the elution buffer. The DNA was quantified with a Nanodrop® ND-1000 spectrophotometer at 260nm. 12µL each amplicon containing 18ng/100basepairs was prepared for sequencing at Microsynth Seqlab GmbH, Göttingen. The sequences obtained were evaluated using the Geneious software (version 5.5.9.0) and for the purpose of comparison, the sequences were compared with NCBI Nucleotide Blast database at (https://blast.ncbi.nlm.nih.gov/Blast.cgi) and TriTryp at (https://tritrypdb.org/tritrypdb/).

### Statistical analysis

We processed our data using Microsoft Excel software and transported it to Graph Pad Prism for the statistical analysis. The trypanosome distribution and infection rate in tsetse flies and pigs within the communities were compared using Chi-squared test. The differences of infection rates were checked between male and female tsetse flies, age groups of pigs, and amongst communities. The comparisons were considered as significantly different at P-value <0.05.

## Results

The overall average apparent density of tsetse flies during the survey period was 4.3 flies/trap/day. We collected over 245 flies but could only dissect 229 comprising 74 males and 155 females respectively (Fig 2). Amongst the communities we visited, tsetse fly infestation was low at Whanabenya while Adenta recorded the highest of the fly numbers (Table 2).

### Molecular detection and identification of trypanosomes in the tsetse fly gut

Nested PCR was performed with DNA extracted from tsetse fly gut tissue and pig whole blood using generic primers to amplify the ITS1 region including part of the 5.8S and 18S genes of *Trypanosomatidae*. Interestingly, trypanosomal DNA was detected in a number of flies from most of the communities, with exception of Ayekokoso (Table 3) with most the trypanosome species almost evenly distributed in tsetse flies between study areas There was simultaneous infection of trypanosomes in the tsetse midgut [32].

Table 2. The rate of occurrence of trypanosome species in tsetse flies and pigs collected in the field

Table 3. Number of trypanosomal DNA detected in the tsetse flies and pigs collected in the study area

The tsetse flies we collected in the field were observed to carry different trypanosome parasites in the midgut as well as different trypanosomes were identified in the blood samples from pigs. The frequency of trypanosomal DNA in tsetse flies was proportional to an increased rate of infection in pigs at the same collection site (figure 3).

**Figure 3.**
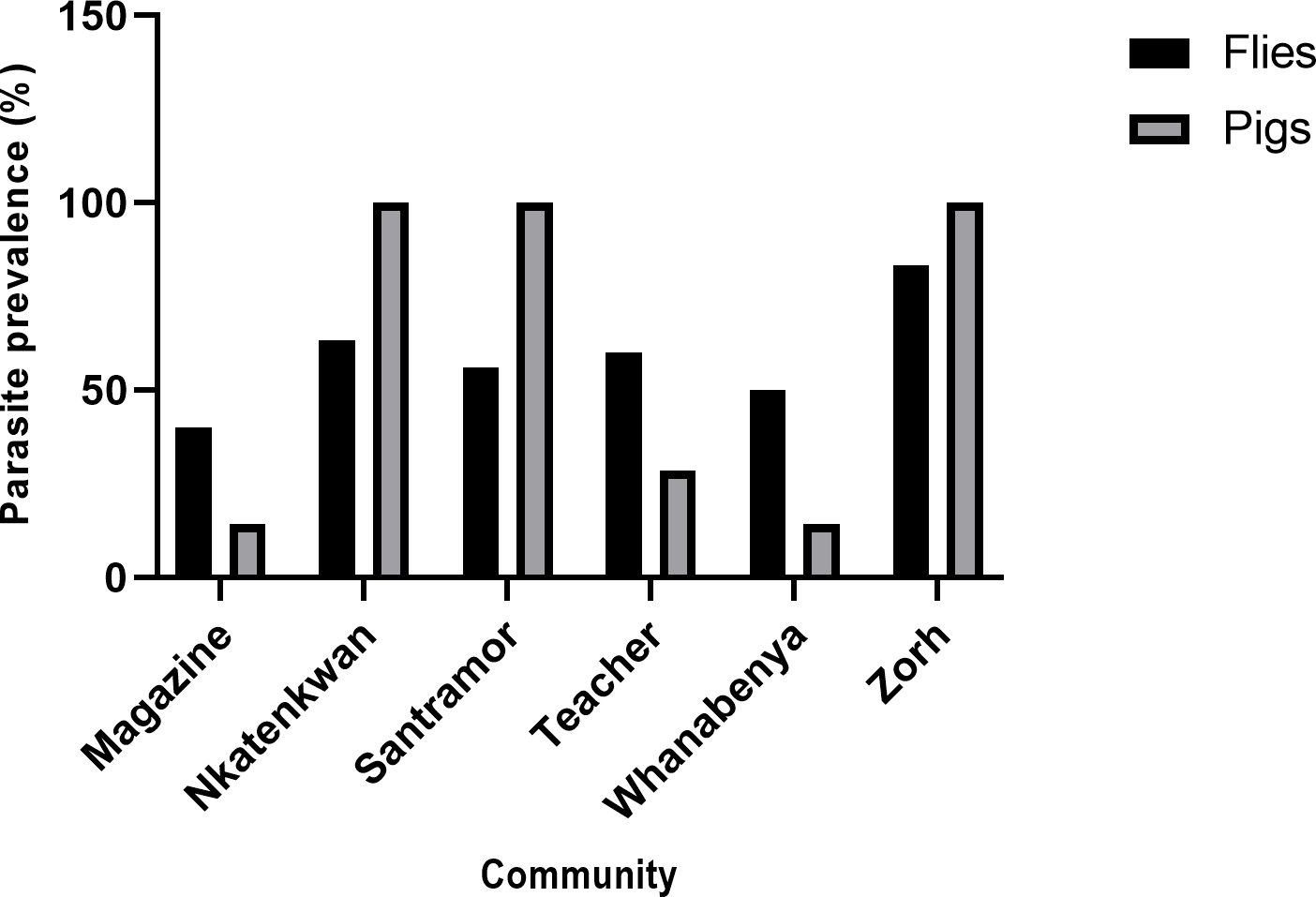
Relationship of the parasite prevalence between tsetse flies and pigs collected from the same communities

**Figure 4.**
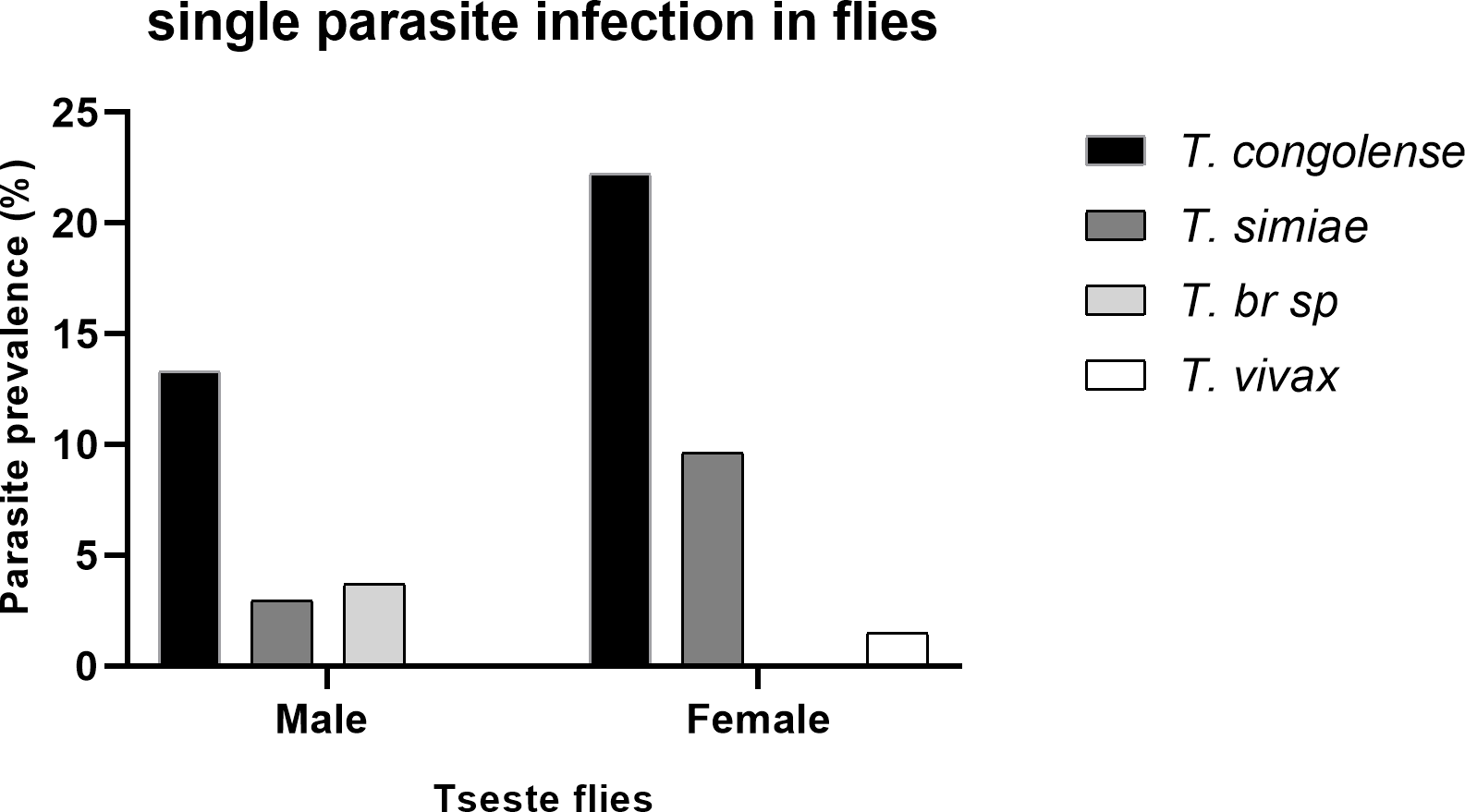
Single and multiple trypanosome infection in collected tsetse flies in relation to fly sex

**Figure 5.**
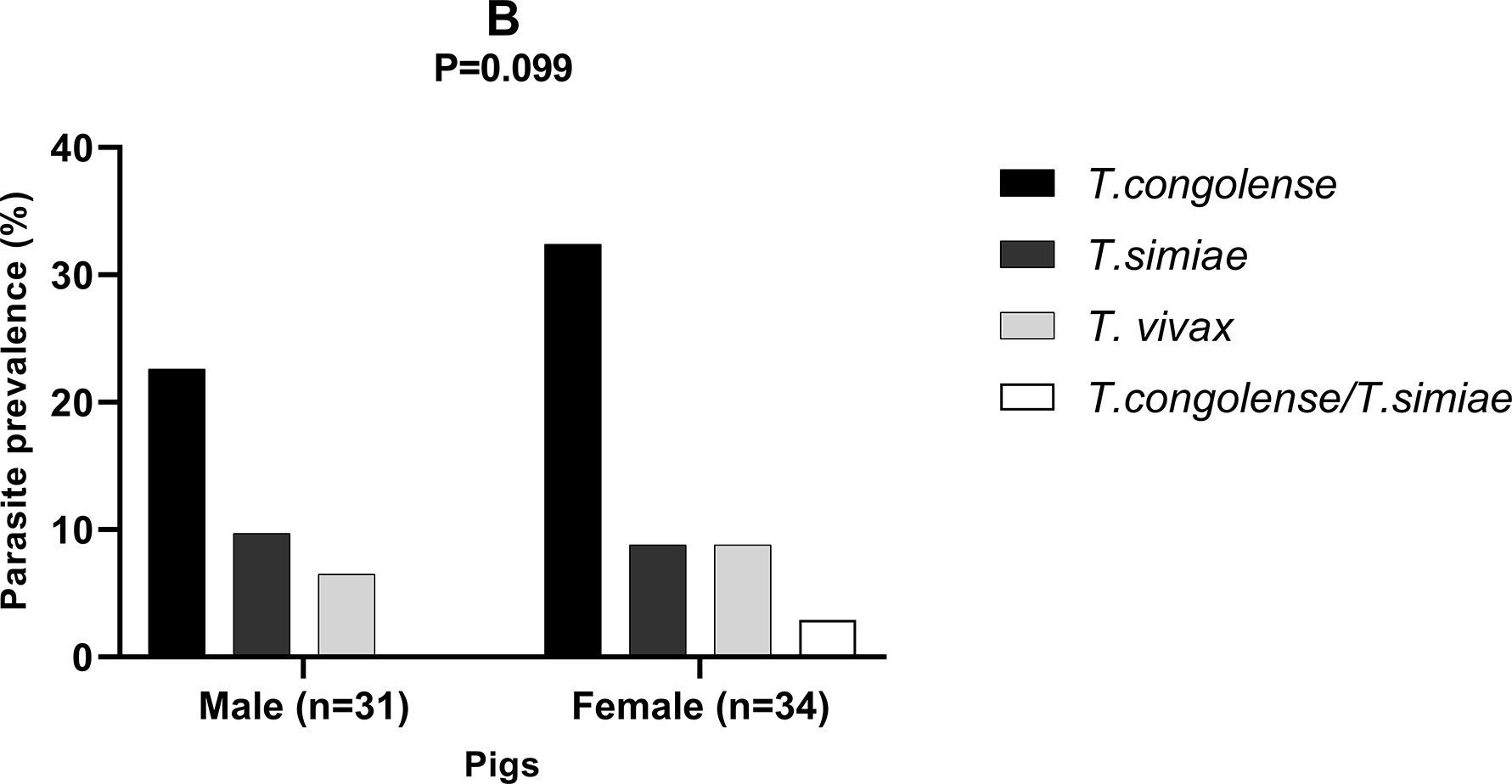
Trypanosome infection within sex and age groups of collected pigs.

**Figure 6.**
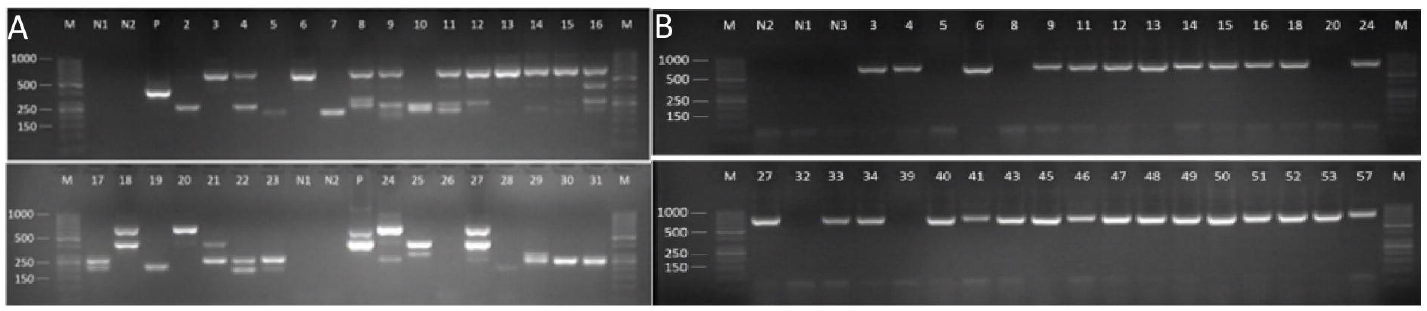
G-stained 2% agarose gel showing amplification of PCR products from tsetse midgut infected with trypanosomes. **A**: Lane (M) molecular marker GeneRuler 50bp Ladder (Thermo Scientific). (N1 and N2): negative controls (from both the first and second reactions), P: positive control (gDNA of *T. brucei*). Amplicons with sizes of ∼700 indicate *T. congolense*, 420-500bp derive from *T. brucei*-like species defined by their similar ITS1 length in this study, the base pairs around 370-400 corresponds to *T. simiae*. Amplicons size of 200-320bp is *T. vivax*. **B**: shows amplification of *T. congolense* (∼700bp fragment size) from *T. congolense*-specific PCR. Lane (M) molecular marker GeneRuler 50bp Ladder (Thermo Scientific). (N1, N2 and N3 are all negative controls: lane 3 to 18 were all *T. congolense* positive samples.)

### *T. congolense*-specific nested PCR

All tsetse fly and pig samples that were considered PCR positive for *T. congolense* were further subjected to species-specific nested PCR to confirm the presence of *T. congolense* and also determine the specific subspecies of *T. congolense* responsible for the infection.). Out of the 48 samples that showed positive for *T. congolense* based on generic ITS1 PCR, 46 of them gave a band size of 760 indicating the presence of forest type and 2 samples did not show any band.

### Molecular detection and identification of trypanosome in cattle and pigs

A total of 20 cattle and 65 pigs from the selected communities were screened for presence of trypanosomes by nested PCR targeting ITS1. None of the cattle was found positive for trypanosomal DNA whereas almost half the number of the pigs screened harboured trypanosomal DNA (Table 2): 46.2% (95% CI: 34.6–58.1) out of 65 pigs were positive for different trypanosome species. The predominant trypanosome species circulating in pigs was *T. congolense* (forest type) and *T. simiae*. The trypanosome infection rate was highest with 100% each at Nkatekwan, Santramor and Zorh. There was significant difference of trypanosome prevalence amongst communities (χ2 = 31.7 95% CI: 34.6–58.1, P=0.0002). The pigs were grouped into three age categories; (i.e. < 1 year, 1-2years and >2years). We encountered only two pigs that were older than two years and they were both positive for trypanosome parasite. 29.2% (95 % CI: 19.6–41.1) of those aged between 1 and 2 years harboured different species of the parasite while 12.3% (95% CI: 6.4–22.5) out of 33 young pigs (i.e.<1year old) were found positive for the parasite.

### Molecular identification of tsetse fly blood meal source

*Cytochrome b gene* DNA was amplified from 10 randomly selected tsetse blood meals in order to identify the blood host. They were purified and prepared for sequencing. Five (5) out of the ten (10) samples gave readable DNA sequence while the other 5 failed to produce readable sequences BLAST analysis of the 5 obtained DNA sequences showed the common warthog (1) and pigs (4) as blood meal source (Table 3).

Table 4. Blood meal source determined by comparison with sequences of species in the NCBI GeneBank database.

## Discussion

Trypanosome detection in tsetse flies, cattle and pigs revealed a high diversity and predominance of *T. congolense* and *T. simiae* in the study region.

The study identified the presence of *Glossina palpalis palpalis* in the forest zone of the eastern region and corroborates the report by Ebhodaghe and others [33], in a study conducted to investigate the distribution of tsetse fly in the same region. Tsestse species have varying ecological requirement in order to thrive in a particular environment [27, 28]. For instance, *Glossina palpalis* prefers forest in which desired environmental factors are invariable [29, 30]. However, other species such as *G. mositant* and *G. tachnoides* that have been previousely reported in the northern Ghana were absent in this study. According to the tsetse distribution models, those species usually thrive in a vegetation that is partially altered by human activities [28, 29]. Thus, the absence of *morsitans* and *tachninoides* in the southern part of the country as reported in this study and others could be an indication of a predisposition towards a northerly retreat of these species. All tsetse species can transmit trypanosomes, however, the *palpalis* group is known as the most significant transmitting vector for Trypanosomiasis [40]. They are capable of transmitting *Trypanosoma brucei gambiense*, which causes sleeping sickness in both western and central Africa [41]. They are also efficient in transmitting animal pathogenic trypanosomes including *T. vivax* and *T. congolense*. Thus, their presence in the area poses a challenge to the public health since both the people and livestock are exposed to the bite of the tsetse fly [18]. Apparent densities did not differ significantly amongst the tsetse biotopes as the factors that determine tsetse density including the vegetation cover and the availability of hosts were almost the same for all sites. The overall apparent density recorded was 4.3 flies/trap/day and was slightly higher than that reported in the new-Juaben district in the same region [8]. The possible reasons could be the low density of livestock population, and/or the use of un-baited traps for the collection of the flies.

### Trypanosome infection in tsetse fly

More than half of the investigated tsetse flies were found carrying trypanosomal DNA in the midgut. Whenever tsetse fly takes up infected blood meal, several trypanosome parasites are mostly cleared away from the midgut by the tsetse immune system. Thus, a few trypanosomes are able to escape the defence mechanism of the tsetse immunity. For these reasons, many trypanosomes are unable to mature and migrate from the midgut to the salivary gland. This could probably be the reason for the high percentage of trypanosomal DNA in the tsetse midgut. It is good to note that DNA is relatively stable and can circulate in body tissues over a period of time and PCR is relatively a sensitive molecular tool to detect and identify even residual traces of trypanosome DNA in varying hosts [45]. Therefore, the percentages shown here are most likely not all reflecting ongoing infections and thus potentially infective flies as the method cannot differentiate between an ongoing infection and an already cleared infections.

Therefore, the presence of DNA alone in a host may not be enough justification for the presence of the active trypanosome parasite

As mentioned earlier, more female tsetse flies collected in this study were found positive for trypanosomal DNA than in the male flies (Fig 3). The lower prevalence of trypanosome in the male tsetse flies does not agree with the literature as few studies have assessed the influence of sex on susceptibility of trypanosome. It appeared the male tsetse was more susceptible than female tsetse in the following studies [36, 37]. The reason could be due to the uneven sex ratios of tsetse flies captured in this study which probably skewed the statistical tendency towards the female flies with the prevalence 71%, of *T. congolense*. compared with the prevalence 40%, of *T. congolense* in males. *T. congolense* was the dominant trypanosome species in the study area with the prevalence of 45.8%. This was higher than that reported in a study carried out in the same region [17]. Ngomtcho and others found a much lower infection rate (24.8%) of *T. congolense* in the same vector species *G. palpalis palpalis* in northern Cameroon. There was an evidence of concurrent colonisation of mixed trypanosome infection in the tsetse midgut. This might come from tsetse blood meal infected with different trypanosome species or successive blood meal uptake from different pigs with different trypanosomes species.

### Trypanosoma species in pigs and cattle

This study revealed trypanosome infection rate of 46.1%, and 0% in pigs and cattle respectively. As only 20 cattle were investigated, the absence of trypanomal DNA in cattle might be explained by the low sample size.

For pigs, however, the infection rate was higher than that reported by Baure et al., in a study carried out in the same region [19]. The trypanosome infection in livestock as revealed in the current study is in line with report by other authors [38–43] and this suggests that pigs are major reservoirs of trypanosomiasis as reported in Cameroon [44–46]. This also implies that other domesticated animals such as dogs and goats might be harbouring the parasites in the area as reported in other parts of Africa [54, 55, 42]. In this study, three trypanosome species were identified with *T. congolense* as the predominant species in the pigs and this corroborate the findings from [47–48], while about 20% and 16% were found carrying *T. simiae* and *T. vivax* respectively. The occurrence of *T*.*vivax* in pigs as found in this study is consistent with the results found in a study by Penchenier and others [59] where species–specific PCR was performed to reveal high levels of *T. vivax* infection in pigs. Interestingly, we observed that high trypanosome prevalence in tsetse flies corresponded to an increased in trypanosomal DNA in pigs from the same sampling site (Fig 3).

We screened multiple cattle breeds including ‘Zebu’, and ‘Ndama’ that are generally known to be trypanotolerant and one susceptible breed, Sanga [60]. Interestingly, no DNA was found in any of the cattle, although both *T. congolense* and *T. vivax* are effectively infecting cattle. This could be attributed to the regular use of trypanocides in even the susceptible breed. This is consistent with the report in a study conducted to investigate trypanosomes in cattle in southern Ghana [61]. We observed low tsetse densities in some of the sampling areas including the cattle ranch site although the frequent administration of trypanocides cannot be excluded. The findings in this study could be attributed to the control practices ensured by the cattle farmers as they asserted that occasionally veterinary officers are invited to administer trypanocidal medication to the cattle. Also, the farmers occasionally spray the cattle against all biting flies which might have reduced the tsetse fly-cattle contact on the farms.

### Blood meal analysis

Sequence analysis of the cytochrome oxidase b gene revealed two vertebrates; *Sus scrofa domesticus* (domestic pig) and *Phacochoerus africanus* (warthog) as host species (Table.3). The availability of blood meal source is one of the most important factors for tsetse fly to thrive in a habitat [62].Thus, the presence of vertebrate host including pigs are key indicator of the presence of tsetse fly in an area. For that reason, the sampling strategy was skewed towards blood meal source (pigs). Nevertheless, tsetse flies are known to take blood meal from a wide range of vertebrate host and can even adopt to feeding from new host in the absence of the usual host [63]. Subsequently, all but one fly had fed from domestic pigs. This fly that revealed the DNA of *Phacochoerus africanus* (warthog) was collected from Nkatekwan. This community was surrounded by secondary forest that could serves as habitat for both the tsetse flies and the *Phacochoerus africanus*.

The outcome of the current study revealed that *G. palpalis palpalis* is the only tsetse fly species in the Suhum/Kraboa/Coaltar district responsible for the transmission of trypanosomes in pigs. ITS1 PCR technique shows that *T. congolense, T. brucei*, and *T. simiae* infection were present in the tsetse fly with *Trypanosoma congolense* being the predominant trypanosome species in the study area. Therefore, scheming an environmentally-friendly control approaches to reduce both the reservoir host and the tsetse fly vector is needed to reduce the prevalence of nagana in the Suhum/Kraboa/Coaltar district. Also, further studies involving a wide range of domestic animals should be carried out to determine prevalence and risk factors of nagana in different seasons to establish seasonal variation in the animal trypanosomiasis transmission.

## Acknowledgement

We would like to give thanks to the Department of Theoretical and Applied Biology (KNUST), Kumasi Centre for Collaborative Research (KCCR), Department of Biology and Chemistry (University of Bremen), Vector-Born Infectious Disease Group (VBID-G), Kroaboa/Coaltar and the Ayensuano districts health directorate for their support. We are particularly grateful to Mr Gbeve and all the farmers who were part of this study for Technical assistance.

