## Supplementary material for "The transmission of African Animal Trypanosomiasis (Nagana) in two districts in the forest zone of Ghana": Tables

| Primer | Direction of the sequence (5’ 3’) | Target | Tm | Expected band size |
| --- | --- | --- | --- | --- |
| ITS OutF  ITS OutR  ITS1 InF  ITS1 InR | TGC AAT TAT TGG TCG CGC  CTT TGC TGC GTT CTT  TAG AGG AAG CAA AAG  AAG CCA AGT CAT CCA TCG | All Trypanosome sp. | 54^°^C | variable |
| TcITS1-OutF  TcITS1-OutR  TcITS1-InF  TcITS1-InR | TGCAATTATTGGTCGCGC  TGTTGGTCGACACTGAGA  TCGCGTGTCTCACGT  TCAAAGATTGGGCAATGT | *T. congolense* | 54^°^C/56^°^C | 681(kilifi)/ 781 (forest) |
| COI- F  COI- R | TTGATTTTTTGGTCATCCAGAAGT  TGAAGCTTAAATTCATTGCACTAATC | Generic Glossina | 55^°^C | 900 |
| VF1d_t1  VR1d_t1 | TGTAAAACGACGGCCAGTTCTCAACCAACCACAARGAYATYGG  CAGGAAACAGCTATGACTAGACTTCTGGGTGGCCRAARAAYCA | All vertebrates | 55°C | 500 |

**Table 1.** Primers, annealing temperature and target as used in this study

Abbreviations: *T.con*, *Trypanosoma congolense*; TGR, *Trypanosoma grayi*;

COI, cytochrome c oxidase 1; Out, outer primer; In, inner primer; F, forward; R, reverse; Tm, annealing temperature

**Table 2.** The prevalence of trypanosome species in tsetse flies and pigs collected in the field

| Trypanosome species | Frequency (%) | |
| --- | --- | --- |
|  | **Tsetse flies** | **Pigs** |
| *T. congolense / T.simiae* | 38.52 | 3.33 |
| *T. simiae* | 14.07 | 20.00 |
| *T. congolense* | 33.33 | 60.00 |
| *T. congolense /T. vivax* | 0.74 | 0.00 |
| *T. congolense /T. brucei. species* | 7.41 | 0.00 |
| *T. brucei. species* | 3.70 | 0.00 |
| *T. vivax* | 1.48 | 16.00 |

Table 3. Number of trypanosomal DNA detected in the tsetse flies and pigs collected in the study area

| **FLIES** | **Total No. Examined** | **PCR Results (+)** | **Prevalence%** | **(95% CI)** | **X^2^** | **P-value** |
| --- | --- | --- | --- | --- | --- | --- |
| Adenta | 71 | 45 | 63.4 | 51.8-73.6 |  |  |
| Ayekokoso | 8 | 0 | 0.0 | 0.0-32.4 |  |  |
| Magazine | 10 | 4 | 40.0 | 16.8-68.2 |  |  |
| Nkatenkwan | 49 | 31 | 63.3 | 49.3-68.2 |  |  |
| Santramor | 57 | 32 | 56.1 | 43.3-68.2 | 16.42 | 0.0367 |
| Teacher | 5 | 3 | 60.0 | 23.1-92.9 |  |  |
| Whanabenya | 4 | 2 | 50.0 | 8.9-91.1 |  |  |
| Pinto | 19 | 13 | 68.5 | 46.0-84.6 |  |  |
| Zorh | 6 | 5 | 83.3 | 43.7-99.2 |  |  |
| **PIGS** |  |  |  |  |  |  |
| Magazine | 7 | 1 | 14.3 | 0.7-51.3 |  |  |
| Nkatenkwan | 10 | 10 | 100.0 | 72.3-100.0 |  |  |
| Santramor | 4 | 4 | 100.0 | 51.0-100.0 |  |  |
| Teacher | 7 | 2 | 28.6 | 5.1-64.1 |  |  |
| Whanabenya | 7 | 1 | 14.3 | 0.7-51.3 |  |  |
| Zorh | 5 | 3 | 60.0 | 23.1-92.9 | 28.06 | 0.0009 |
| Bonkua No.1 | 7 | 2 | 28.6 | 5.1-64.1 |  |  |
| Budu | 7 | 1 | 14.3 | 0.7-51.3 |  |  |
| Nankese | 2 | 2 | 100.0 | 17.8-100.0 |  |  |
| Mensakrom | 7 | 2 | 28.6 | 5.1-64.1 |  |  |
| **CATTLE** |  |  |  |  |  |  |
| Ayekokooso | 10 | 0 | 0.0 | 0.0-27.8 |  |  |
| Akorabo | 10 | 0 | 0.0 | 0.0-27.8 |  |  |

χ2: Chi-square test;	**highly significant, CI: Confidence Interval

**Table 4.** Blood meal source determined by comparison with sequences of species in the NCBI GeneBank database.

| Mammalian host species | Cyt b detection rate | NCBI (Genebank accession number) |
| --- | --- | --- |
| *Phacochoerus africanus* (Common warthog) | 1/28 | KJ192881 |
| *Sus scrofa breed Fengjing mitochondrion* (Pig) | 4/28 | MH603005.1 |
